# Adipose Tissue Analysis Toolkit (ATAT) for Automated Analysis of Adipocyte Size and Extracellular Matrix in White Adipose Tissue

**DOI:** 10.1101/2023.12.12.571339

**Authors:** Jacob J. Robino, Alexander P. Plekhanov, Qingzhang Zhu, Michael D. Jensen, Philipp E. Scherer, Charles T. Roberts, Oleg Varlamov

## Abstract

**Objective:** The pathological expansion of white adipose tissue (WAT) in obesity involves adipocyte hypertrophy accompanied by expansion of collagen-rich pericellular extracellular matrix (ECM) and the development of crown-like structures (CLS). Traditionally, WAT morphology is assessed through immunohistochemical analysis of WAT sections. However, manual analysis of large histological sections is time-consuming, and available digital tools for analyzing adipocyte size and pericellular ECM are limited. To address this gap, we developed the Adipose Tissue Analysis Toolkit (ATAT), an ImageJ plugin facilitating analysis of adipocyte size, WAT ECM and CLS.

**Methods and Results:** ATAT utilizes local and image-level differentials in pixel intensity to independently threshold background, distinguishing adipocyte-free tissue without user input. It accurately captures adipocytes in histological sections stained with common dyes and automates the analysis of adipocyte cross-sectional area, total-field, and localized region-of-interest ECM. ATAT allows fully automated batch analysis of histological images using default or user-defined adipocyte detection parameters.

**Conclusions:** ATAT provides several advantages over existing WAT image analysis tools, enabling high-throughput analyses of adipocyte-specific parameters and facilitating the assessment of ECM changes associated with WAT remodeling due to weight changes and other pathophysiological alterations that affect WAT function.

**Study Importance Questions:** *What is already known about this subject?:* The manual analysis of large WAT histological sections is very time-consuming, while digital tools for the analysis of WAT are limited.

*What are the new findings in your manuscript?:*

- ATAT enables fully automated analysis of batches of histological images using either default or user-defined adipocyte detection parameters
- ATAT allows high-throughput analyses of adipocyte-specific parameters and pericellular extracellular matrix
- ATAT enables the assessment of fibrotic changes associated with WAT remodeling and crown-like structures

*How might your results change the direction of research or the focus of clinical practice?:*

- ATAT is designed to work with histological sections and digital images obtained using a slide scanner or a microscope.
- This tool will help basic and clinical researchers to conduct automated analyses of adipose tissue histological sections.

## Introduction

Collagen is the major structural component of healthy white adipose tissue (WAT) that organizes the extracellular matrix (ECM) into three-dimensional networks that control adipocyte function (1). ECM surrounding individual adipocytes, rather than total ECM deposition, is elevated in WAT of obese individuals (1, 2, 3, 4, 5). In lean healthy individuals, pericellular ECM is minimal, being present only in collagen-rich septa that compartmentalize WAT into smaller lobes (3, 6). WAT becomes less lobular in response to diet-induced obesity, enabling WAT to expand in response to adipocyte hypertrophy (6). Pathological expansion of ECM can also lead to the accumulation of proinflammatory immune cells (2, 7, 8, 9). The area of the tissue occupied by ECM may change dynamically in response to changing physiological conditions. For example, WAT expansion can lead to a pathological state characterized, in large part, by the induction of local inflammation and hypoxia, resulting in excess ECM accumulation. Therefore, ECM remodeling may drive adipocyte dysfunction and promote a microenvironment conducive to metabolic dysfunction (2). Quantitative assessment of ECM may, therefore, predict a pathophysiological state of WAT that contributes to disease progression. The recruitment of macrophages and other immune cells to the pericellular ECM to form crown-like structures (CLS) has been shown to contribute to obesity-induced ectopic lipid accumulation (10). These structures are characterized by the presence of CD68+ macrophages surrounding dead adipocytes (11, 12, 13). CLS can undergo dynamic changes in number and cellular composition in response to obesity and weight loss (11, 14).

The primary goal of this study was to develop a computer program capable of automatically capturing adipocyte boundaries and performing calculations for adipocyte size and pericellular ECM thickness. Additionally, we aimed at the analysis of larger fibrotic areas within WAT that are known to be induced by various pathological processes, including tumor invasion (15, 16, 17), obesity (2, 3, 4), and HIV infection (18, 19, 20), as well as the analysis of WAT septa involved in tissue remodeling during weight loss (4). While there are several digital tools available for the analysis of adipocyte size, including AdipoCount, HALO, and Adiposoft (21, 22, 23, 24), there is a paucity of image analysis programs for automated analysis of WAT ECM.

Consequently, a software tailored for pericellular ECM analysis must meet two critical criteria: first, it should be proficient in excluding areas in an image containing other features, including blood vessels; and second, it must incorporate an algorithm for measuring ECM thickness around individual adipocytes. In response to this need, we developed Adipose Tissue Analysis Toolkit (ATAT), a software program that satisfies both of these requirements. ATAT is designed to analyze large batches of images with minimal user input, offering an efficient solution to the automated or user-unassisted image analyses of adipocyte size and pericellular ECM in WAT using the open-source image analysis software ImageJ (22).

## Methods

### Animal studies

This study was approved by the Oregon National Primate Research Center Institutional Animal Care and Use Committee and conforms to current Office of Laboratory Animal Welfare regulations as stipulated in assurance number A3304-01. WAT samples were collected from necropsies of adult female rhesus macaques performed as part of an unrelated study.

### Human studies

Abdominal adipose tissue biopsies were collected as part of studies approved by the Mayo Clinic IRB. Samples were collected under sterile conditions using local anesthesia and processed for immunohistochemistry as previously described (25).

### Macaque tissue processing and staining

WAT histology and image analysis were performed as described (26). Briefly, 200 to 500-mg fragments of subcutaneous (SC)-WAT and omental (OM)-WAT were collected at necropsy and fixed in zinc formalin (Fisher Scientific, Hampton, NH, USA) at room temperature for 48 hours. Samples were transferred to 70% (v/v) ethanol for 5 days, embedded in Paraplast wax (Leica, Wetzlar, Germany), and 5-μm sections were prepared using a micron rotary microtome. Slides were stained with the Picro-Sirius Red Stain Kit (Abcam, Boston, MA, USA) or the NovaUltra^TM^ H&E Stain Kit (IHC World LLC, Ellicott City, MD, USA) according to the manufacturer’s instructions.

### Image capture and analysis

Images representing tissue segments approximately 5 to 10 mm in size were acquired using an Aperio AT2 System slide scanner (Leica Biosystems, Wetzlar, Germany) and saved as TIFF files.

Image analysis was performed using ImageJ and the ATAT plugin as described in the instruction manual. Data analysis was conducted in ImageJ and Excel.

### Online resource

The ATAT plugin, the instruction manual and a source code can be downloaded from latest releases on the GitHub page (https://github.com/aplekh/ATAT).

## Results

### Detection of adipocytes through automatic thresholding

The intricate composition and complex anatomical structure of WAT pose challenges for analysis. In thin paraffin-embedded histological sections, adipocytes are visually represented as cell ghosts containing plasma membranes and delipidated lipid droplets (Figure 1A). The non-adipocyte fraction encompasses a collagen-rich ECM containing immune cells, fibroblasts, stem cells, blood vessels, and nerve fibers. The primary purpose of the ATAT is to distinguish between the background (comprising the interior of adipocytes) and the foreground (encompassing adipocyte membranes and ECM). To achieve this, ATAT identifies the brightest peak in the pixel intensity distribution and scans the histogram leftwards until it locates a peak in the second derivative. Figure 1A depicts an original H&E-stained WAT image. ATAT generates a histogram of the pixel intensity distribution, with the red line indicating the brightest peak corresponding to the modal intensity of the background (Figure 1B). The blue line marks the first peak of the second derivative found by ATAT during the leftward scan. When applied as a threshold, the initial image is successfully thresholded (Figure 1D). Recognizing that pixel intensity distributions can vary across images, ATAT allows the calculation of multiple thresholds. In this example, a 16-region thresholding approach is employed (Figure 1C).

**Figure 1.**
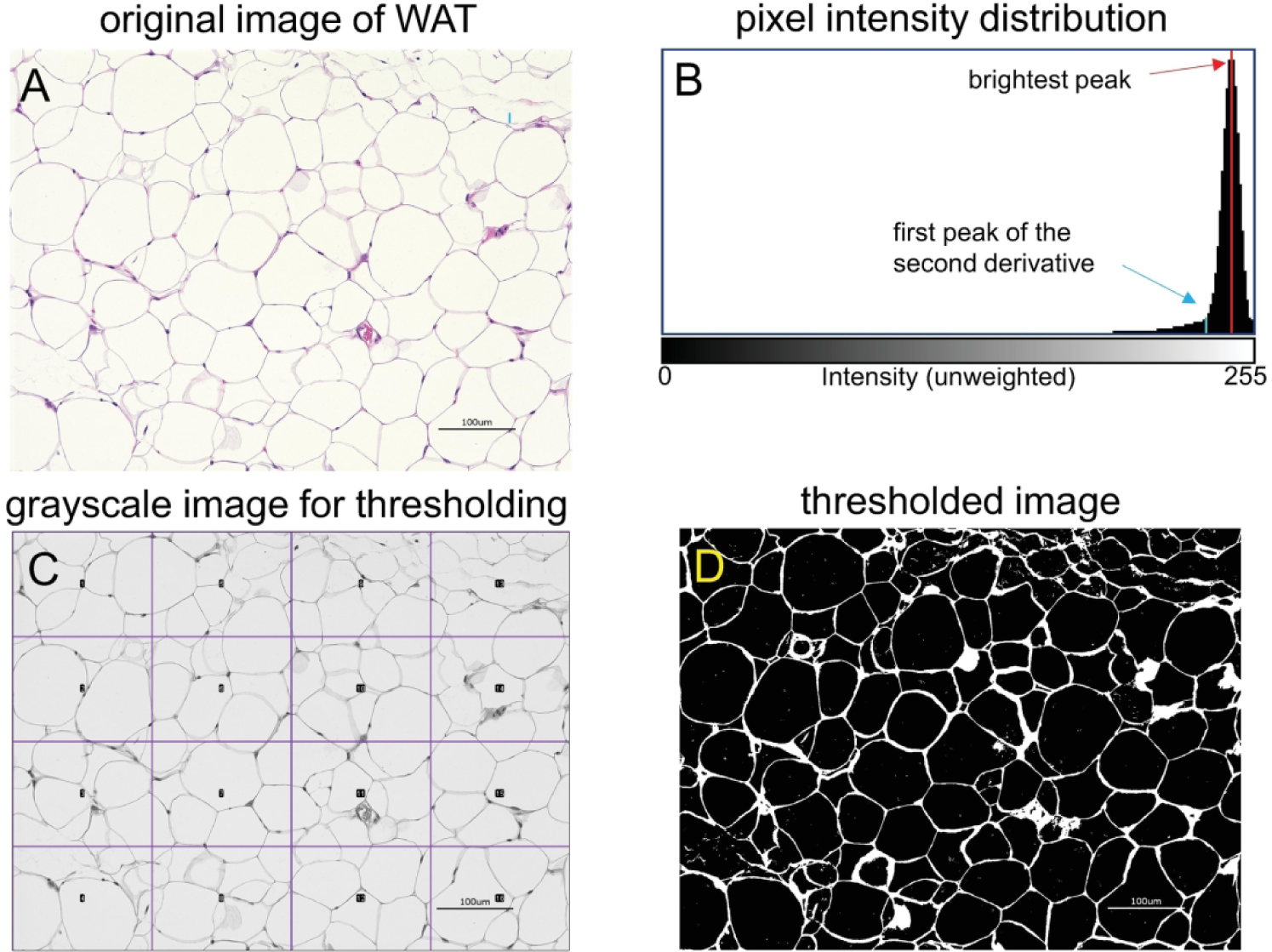
Automatic thresholding. (A) Original H&E-stained image of macaque OM-WAT. (B) Histogram of the pixel intensity distribution; the red line indicates the brightest peak of the distribution, equal to the modal intensity of the image’s background. The blue line indicates the first peak of the second derivative that ATAT finds as it scans leftwards; when applied as a threshold, an image (A) is successfully thresholded (D). (C) ATAT allows multiple thresholds to be calculated and applied to each image. A 16-region thresholding approach is used in this image. Scale bar, 100 μm.

### Adipocyte boundary detection and adipocyte size analysis

This feature is designed to specifically capture adipocytes and calculate their cross-sectional areas. ATAT requires user input to set specific adipocyte parameters, which include the minimal and maximal adipocyte size (area) and circularity. Figure 2 visually demonstrates the detection of adipocyte boundaries in a subcutaneous WAT (SC-WAT) sample from a SIV-infected female rhesus macaque. Figure 2A depicts a representative WAT sample stained with Picro-Sirius Red, revealing prominent adipocyte-free fibrotic areas encircled by adipocyte-rich regions. Figure 2B provides an enlarged view of an area displaying adipocytes adjacent to a region containing substantial fibrosis. ATAT is designed to automatically exclude large fibrotic areas and selectively capture adipocytes, streamlining the analysis process. Figure 2C illustrates the adipocyte capture image generated by ATAT.

**Figure 2.**
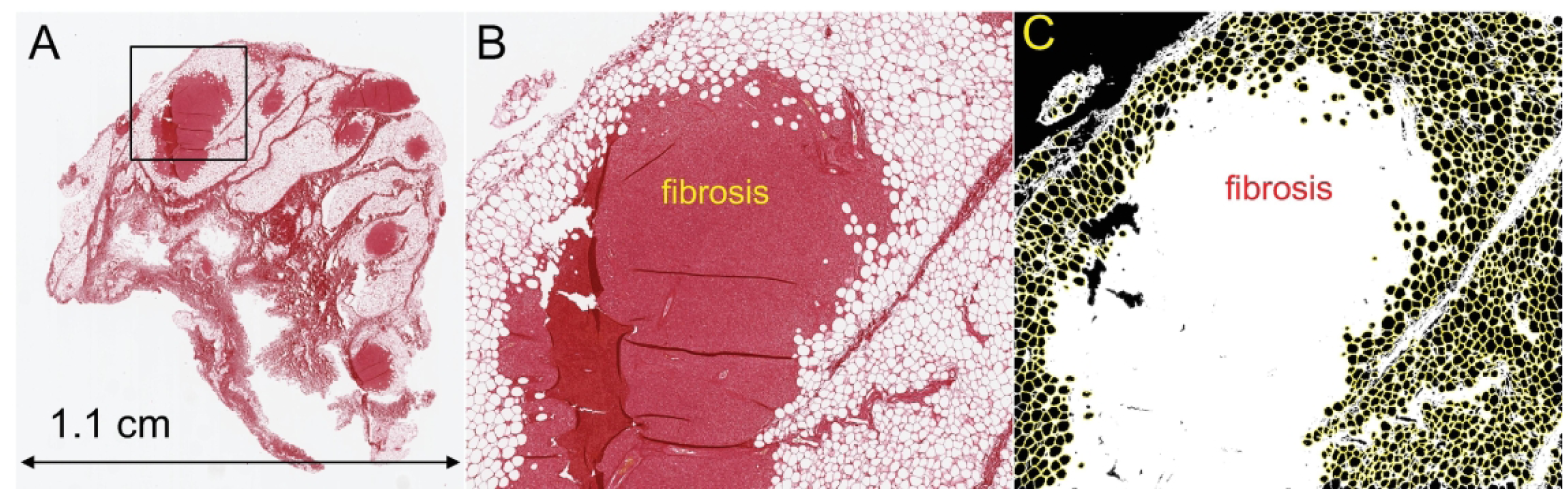
Adipocyte boundary detection. (A) A representative image of SC-WAT of an SIV-infected rhesus macaque stained with Picro-Sirius Red. Tissue fibrosis is stained red. (B) Enlarged area of WAT showing adipocytes adjacent to the fibrotic area. (C) Adipocyte capture performed by ATAT. Digital images were thresholded, identifying all foreground pixels as ECM or fibrotic areas. Background pixels were used to identify adipocytes and characterize their morphology, including sectional area. Adipocyte capture was conducted with the ImageJ “Analyze Particles” command, with minimum and maximum particle sizes set to 300 and 30,000 square microns, respectively, and with a minimum circularity of 0.4.

### Comparing ATAT and Adiposoft for adipocyte detection

We conducted a comparative analysis of ATAT and a commercially available software, Adiposoft, for the assessment of adipocyte size in histological sections. Both software programs facilitate whole-image analysis of WAT, capturing individual adipocytes and measuring their areas (Supplementary Figure S1A). Notably, in contrast to Adiposoft. ATAT demonstrates superior accuracy in capturing adipocyte boundaries by delineating the region of interest (ROI) closer to the plasma membrane (Supplementary Figure S1B), resulting in slightly larger adipocyte areas (Supplementary Figures S1D and E). Additionally, ATAT effectively excludes potential artifacts involving ruptured adipocyte membranes, while Adiposoft occasionally identifies larger objects comprising fused adipocytes (Supplementary Figure S1C). An important feature of ATAT is its capability for region-specific analysis of WAT, enabling users to define ROIs that exclude other histological structures such as fibrotic areas and CLS, as described below. In contrast, Adiposoft is limited to whole-image analysis.

### The correlation between ECM thickness and Picro-Sirius Red staining intensity

ATAT performs spatial analysis of the pericellular ECM but does not take into account the density of collagen fibers that compose the ECM. Consequently, our aim was to determine whether the packing density of collagen fibers is associated with the size of pericellular ECM surrounding individual adipocytes. Figure S2A illustrates the analysis of pericellular ECM profiles in the OM-WAT section stained with Picro-Sirius Red. This analysis assesses both the linear size of pericellular ECM and the grey values of pixel intensities associated with Picro-Sirius Red staining (Figure S2B). The analysis of multiple adipocytes reveals that pericellular ECM thickness correlates with Picro-Sirius Red staining intensity (Figure S2C), suggesting that thicker pericellular ECM regions contain a higher collagen density.

### ATAT analysis of vascularized WAT

ATAT can perform whole-image analysis of adipocyte areas in highly vascularized regions of WAT using H&E staining. It is important to note that ATAT does not detect red blood cells within blood vessels visualized by H&E staining. To ensure accurate analysis, users can configure specific circularity, filter, and particle size parameters to exclude potential irregularly shaped artifacts within blood vessels and the adjacent connective tissue (Supplementary Figure S3). Figure S3 illustrates ATAT’s successful detection of adipocyte boundaries within highly vascularized regions of OM-WAT, while excluding non-adipocyte features within blood vessels and the vascular bed.

### Pericellular ECM analysis

This feature is specifically designed for calculating pericellular ECM in WAT. In the analysis of pericellular ECM, it is essential to outline ROIs that include adipocytes while excluding fibrotic areas. ATAT systematically measures the thickness of ECM surrounding each adipocyte in eight radial directions and computes the mean (average), median, and longest ECM thickness per adipocyte (Figure 3A). The software records the data and generates the ECM capture mask (Figure 3B). Figures 3C and D illustrate the analysis of pericellular ECM using perigonadal WAT from mice exposed to either a control or a high-fat diet (HFD). The average, longest, and median pericellular ECM thickness appear higher in WAT from the HFD-exposed mouse compared to control WAT (Figure 3D). Notably, this analysis is adaptable to alternative staining techniques, such as Picro-Sirius Red and Trichrome.

**Figure 3.**
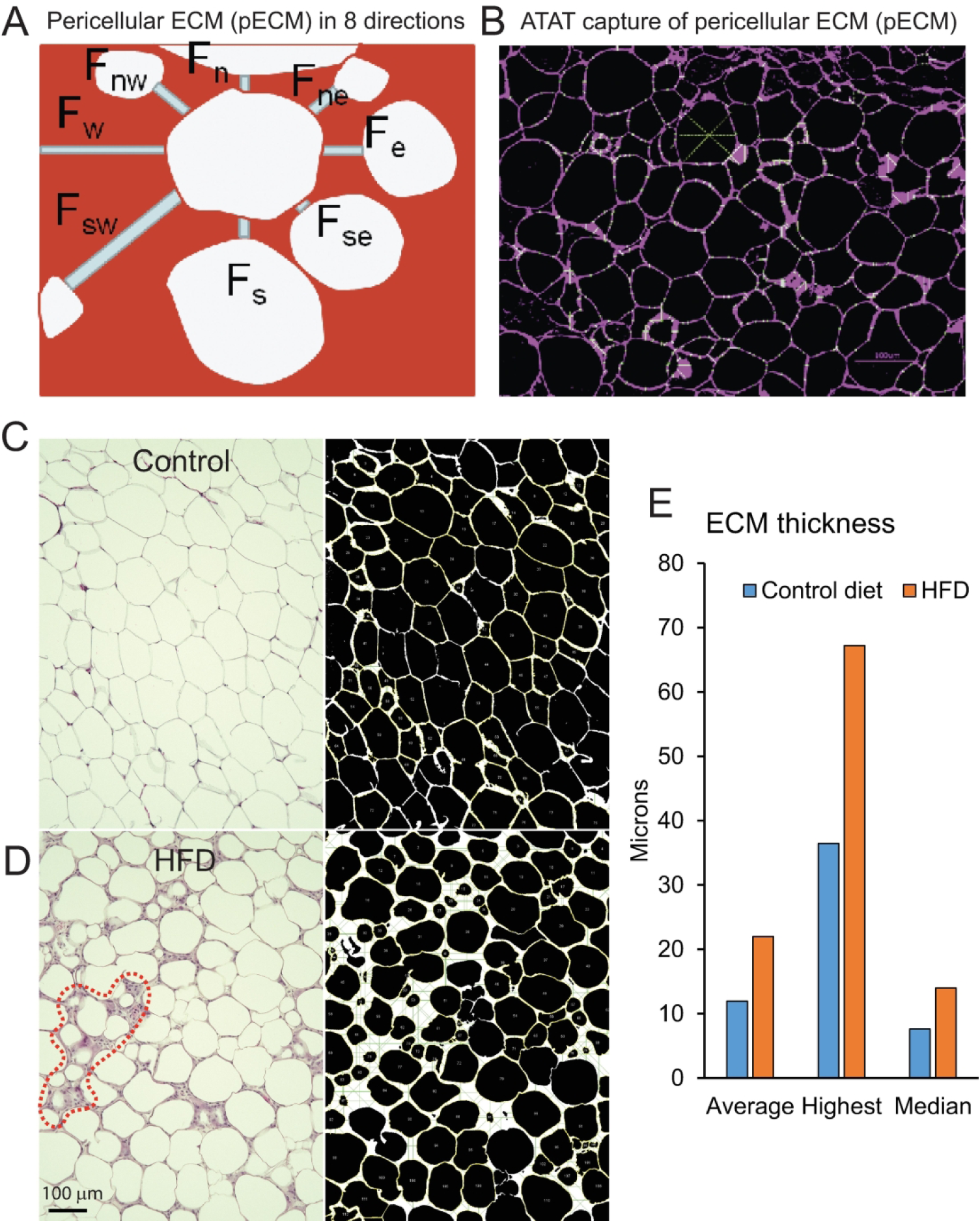
Analysis of pericellular ECM in mouse perigonadal WAT. (A) To quantify WAT pericellular ECM, ATAT measures the thickness of ECM surrounding each adipocyte in eight radial directions and calculates the average, median, and longest ECM thickness per each adipocyte. (B) Thresholded image from WAT sample showing adipocyte boundaries (magenta) and fibrotic lines (green). For illustration purposes, in one of the adipocytes, fibrotic lines project to the center of the cell. (C and D) Representative images of perigonadal WAT from mice exposed to a control diet (C) or high-fat diet (HFD, D). (D) A large crown-like structure is outlined by the red line. Masked images, adipocyte boundaries are outlined by yellow ROIs; green lines, fibrotic lines. (E) The bar graph shows the average, highest, and median ECM thickness for the images in panel C and D (n=133-183 adipocytes).

### Studies of WAT lobular architecture

WAT is subcompartmentalized into millimeter-sized lobes surrounded by sheets of connective tissue that comprise the inter-lobular septa. In two-dimensional histological sections stained with Picro-Sirius Red, the inter-lobular septum appears as a branched belt of connective tissue separating individual WAT lobes (Figure 4A, arrowheads). Employing user-defined analytic ROIs for WAT lobes and septa, ATAT systematically records adipocyte-specific and ECM parameters within each region (Figure 4B). This specific analysis shows that adipocytes located near the septa appear smaller than intra-lobular adipocytes (Figures 4C and D). Additionally, the ECM thickness is notably greater in the septal area compared to the intra-lobular area (Figure 4E).

**Figure 4.**
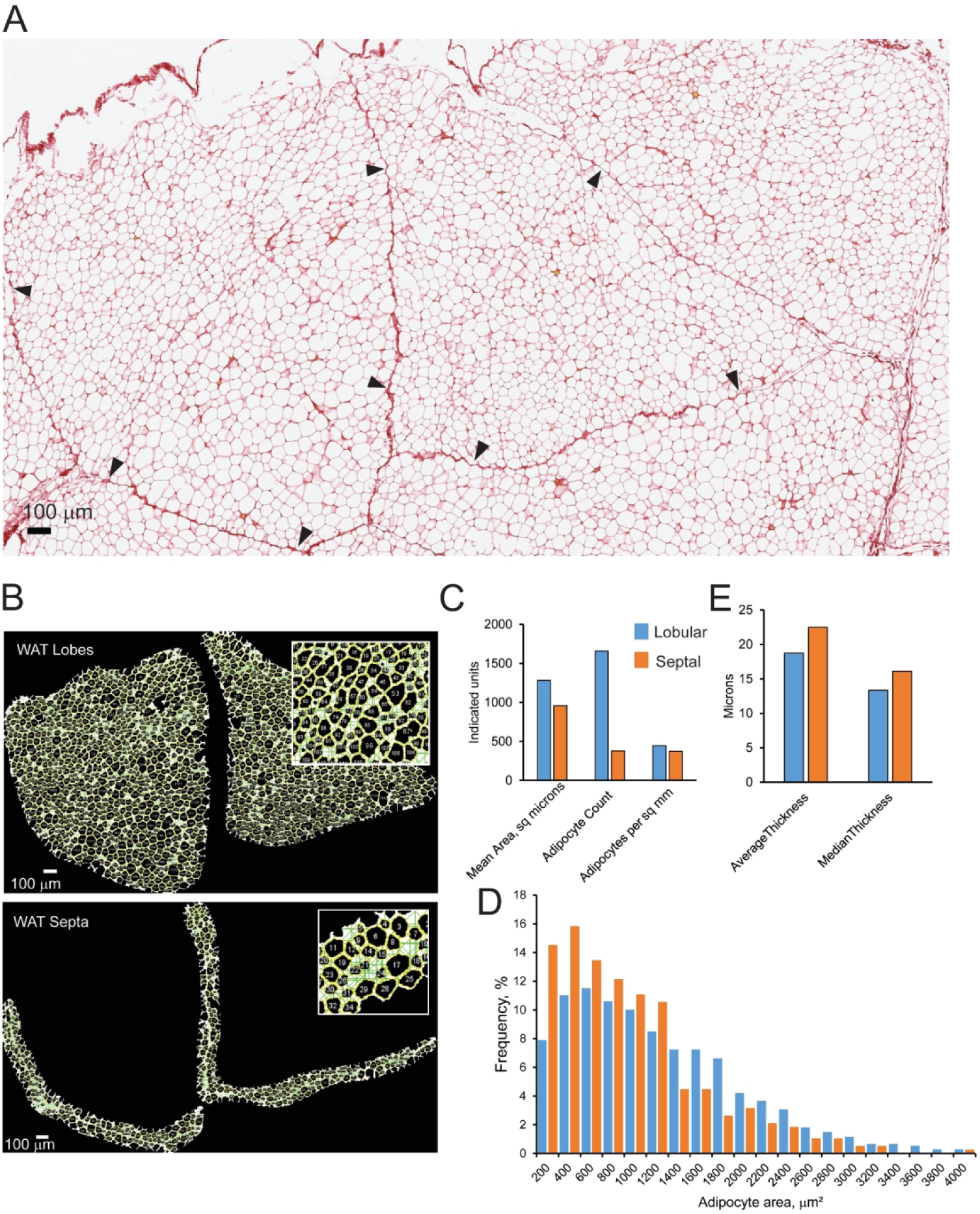
Analysis of lobular structures in WAT. (A) A representative image of macaque OM-WAT showing anatomical lobes separated by the septa (arrowheads) stained with Picro-Sirius Red. (B) ATAT analysis of WAT septa surrounding two lobes; the binary masks show adipocytes (black) and fibrotic lines (green); insets, enlarged masked areas of WAT. (C) Adipocyte parameters calculated for lobular and septal areas of WAT. (D) Histogram of adipocyte size distribution for lobular and septal areas of WAT. (E) ECM parameter calculated for lobular and septal areas of WAT. Bars shown in (C) an (E) are mean values.

### Studies of WAT fibrotic areas

Fibrotic areas, denoting regions within WAT characterized by a disorganized lobular structure (Figure 5A), are discernible on histological sections stained with Picro-Sirius Red. In these fibrotic areas, adipocytes exhibit greater variation in size and shape compared to intralobular adipocytes (Figure 5A, arrowheads). Leveraging user-defined analytic ROIs, ATAT systematically analyzes adipocyte size and ECM parameters in both intra-lobular and fibrotic areas of the image (Figure 5B). A representative analysis illustrates that adipocytes located within fibrotic areas are, on average, smaller than their intra-lobular counterparts (Figures 5C-E). Additionally, ECM thickness is greater in fibrotic areas compared to nonfibrotic regions (Figure 5F).

**Figure 5.**
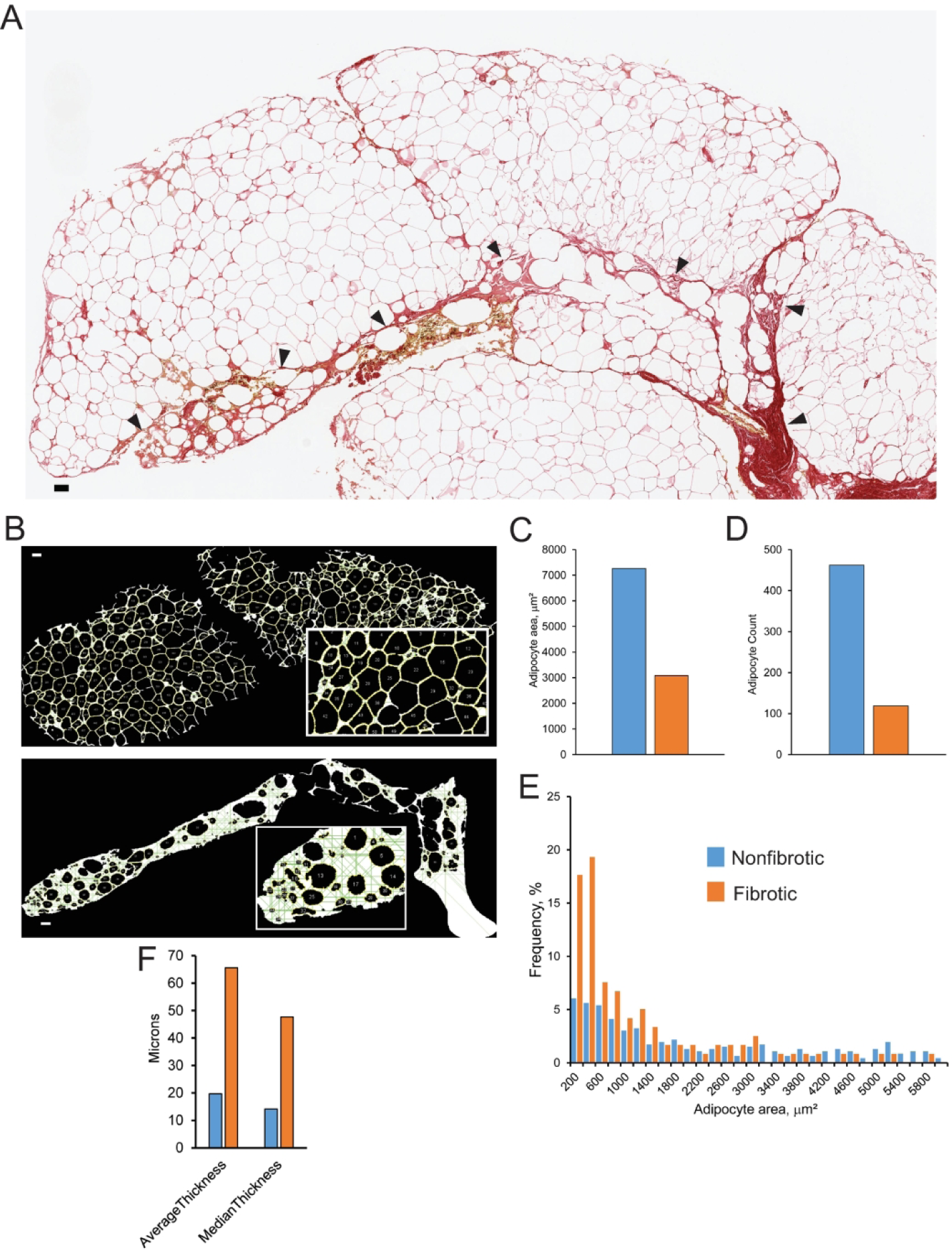
Analysis of WAT fibrotic areas. (A) Representative image of macaque SC-WAT showing normal lobular (nonfibrotic) and fibrotic (arrowheads) areas stained with Picro-Sirius Red, scale bar=100 μm. (B) ATAT analysis of intra-lobular adipocytes (top) and extralobular adipocytes associated with fibrotic areas (bottom); the binary masks show adipocytes (yellow) and fibrotic lines (green); insets, enlarged masked areas of WAT. (C-E) Adipocyte parameters calculated for lobular and fibrotic areas of WAT. (F) ECM parameter calculated for each area of WAT. Bars are mean values; the number of adipocytes (adipocyte count) analyzed in each region is indicated in (D).

### Analysis of CLS

CLS can be visualized in WAT through H&E staining or CD68 immunostaining to show nucleated cells surrounding dying adipocytes (Figures 3D, 6A and 6B). CLS are typically counted manually, but their dimensions are not determined. ATAT facilitates a more in-depth CLS analysis, allowing the evaluation of their dimensions as a potential marker for the spatiotemporal development of a proinflammatory milieu in WAT. Using dedicated modules for ECM analysis, ATAT effectively quantifies CLS dimensions, revealing a thicker ECM associated with CLS compared to normal WAT areas located outside CLS (Figure 6C and E). Dying adipocytes within CLS appear smaller than those outside CLS (Figure 6D). Of particular note, CLS can grow in size to form large CLS, as demonstrated in Figure 3D. Since these large CLS lack defined boundaries, they are not easily countable. In such cases, it is more practical to quantify the average ECM thickness across the larger WAT area instead of focusing on individual CLS.

**Figure 6.**
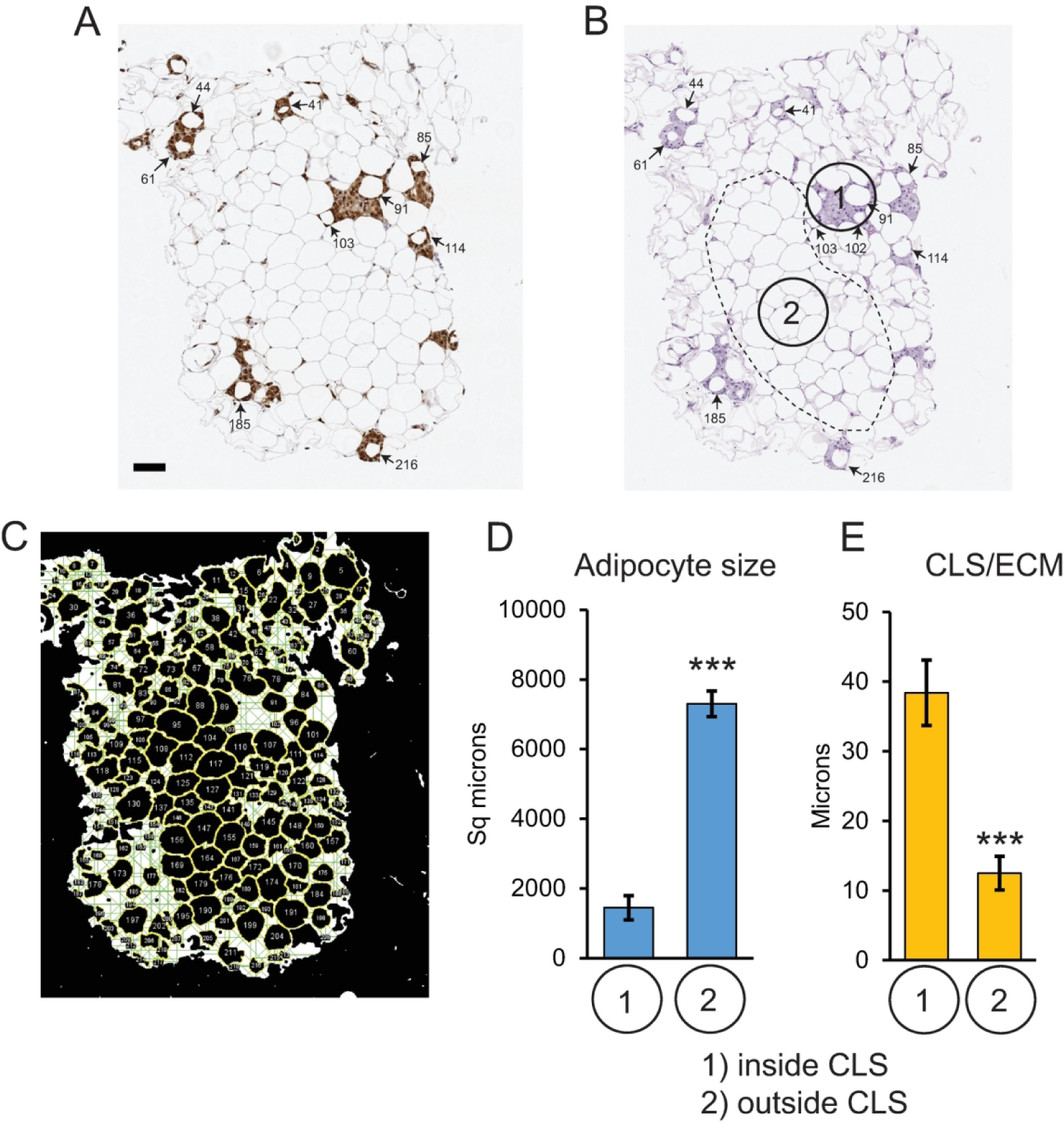
Analysis of CLS in WAT. (A and B) Representative images of human SC-WAT stained with anti-CD68 antibody (A) and H&E (B), depicting macrophage-rich areas of WAT containing smaller adipocytes (arrows and small numbers). The dotted line outlines the macrophage-free stable area of WAT containing larger adipocytes; scale bar=100 μm. (C) ATAT-generated mask of the H&E image showing adipocyte ROIs in yellow and fibrotic lines in green. (D and E) Quantification of average adipocyte area (D) and ECM thickness associated with crown-like structure-positive (1) and crown-like structure-negative (2) areas of WAT. Bars are means ± SEM, n=10 adipocytes per area; T-test, p<0.001.

## Discussion

Disease-associated alterations in WAT morphology are commonly evaluated through histochemical analysis of WAT sections. However, manual analysis of large histological sections of WAT can be excessively time-consuming, while digital tools for the analysis of adipocyte size and pericellular ECM are limited. In response to this need, we have developed the Adipose Tissue Analysis Toolkit (ATAT) for the open-source image analysis software ImageJ. ATAT employs both local and global differentials in pixel intensity to distinguish background apart from foreground without user input. It accurately captures adipocyte areas, cell density, and pericellular ECM thickness in histological sections stained with common histological dyes. The toolkit enables fully automated batch analysis of histological images using default or user-defined adipocyte detection parameters, such as minimum circularity and minimum/maximum cross-sectional area. ATAT’s thresholding techniques exhibit high resistance to false-negative and false-positive detection events compared to other open-source adipocyte analysis software, offering superior measurement accuracy, particularly in low-magnification images. ATAT features several quality-of-life features, including automatic summarization of batch-collected data, and an optional histogram builder. Released as an ImageJ plugin, ATAT has the potential to elevate the rigor and reproducibility of WAT research.

In comparison with other freely available adipocyte quantification tools like Adiposoft (24), AdipoCount (21), and AdipoQ (27), ATAT introduces novel features. While Adiposoft, AdipoCount, and AdipoQ offer automated quantification of adipocyte number and cross-sectional areas in histological sections of WAT, only AdipoQ assesses the intensity of surrounding pixels adjacent to individual adipocytes. Although AdipoQ approximates pericellular ECM, it cannot record the two-dimensional size of the ECM adjacent to each adipocyte. Moreover, AdipoQ’s output may be influenced by staining intensity variations resulting from different histological visualization methods, where darker staining yields greater pixel intensity values. In contrast, ATAT relies on quantifying distances between adjacent adipocytes, offering independence from global and local staining intensity associated with ECM. This approach is largely unaffected by staining variability commonly observed with common histological dyes, making it suitable for batch analysis of images produced by different histological methods, such as H&E, Picro-Sirius Red, trichrome, and 3,3-diaminobenzidine (DAB)-based immunohistochemistry.

### Conclusions

ATAT is designed for use with histological sections and digital images acquired through a slide scanner or a microscope. It offers the flexibility to analyze the entire image or manually-selected ROIs, focusing on adipocytes while excluding other features such as blood vessels and histological staining artefacts. The sequential steps and outcomes of the analysis are illustrated in Figure 7.

**Figure 7.** ATAT analysis pipeline. ATAT is compatible with histological sections stained with H&E, trichrome, or Picro-Sirius Red. Digital images can be acquired using a slide scanner or a microscope. ATAT offers the flexibility to conduct analyses on the entire image or manually selected ROIs that specifically contain adipocytes while excluding other features such as blood vessels and histological staining artifacts. The following sequential steps and results of the analysis are outlined for user guidance.

## Author contribution

JJR developed the initial macro-based algorithms, AP developed the ImageJ plugin and the instruction manual, CTR contributed to the overall study design and wrote the manuscript, MDJ and PES provided samples and reviewed the manuscript, and OV designed the study and wrote the manuscript.

## Acknowledgments

We acknowledge the assistance of the ONPRC Integrated Pathology Core. The AT2 slide scanner used was funded by NIH grant S10 OD025002.

---

**Figure S1.**
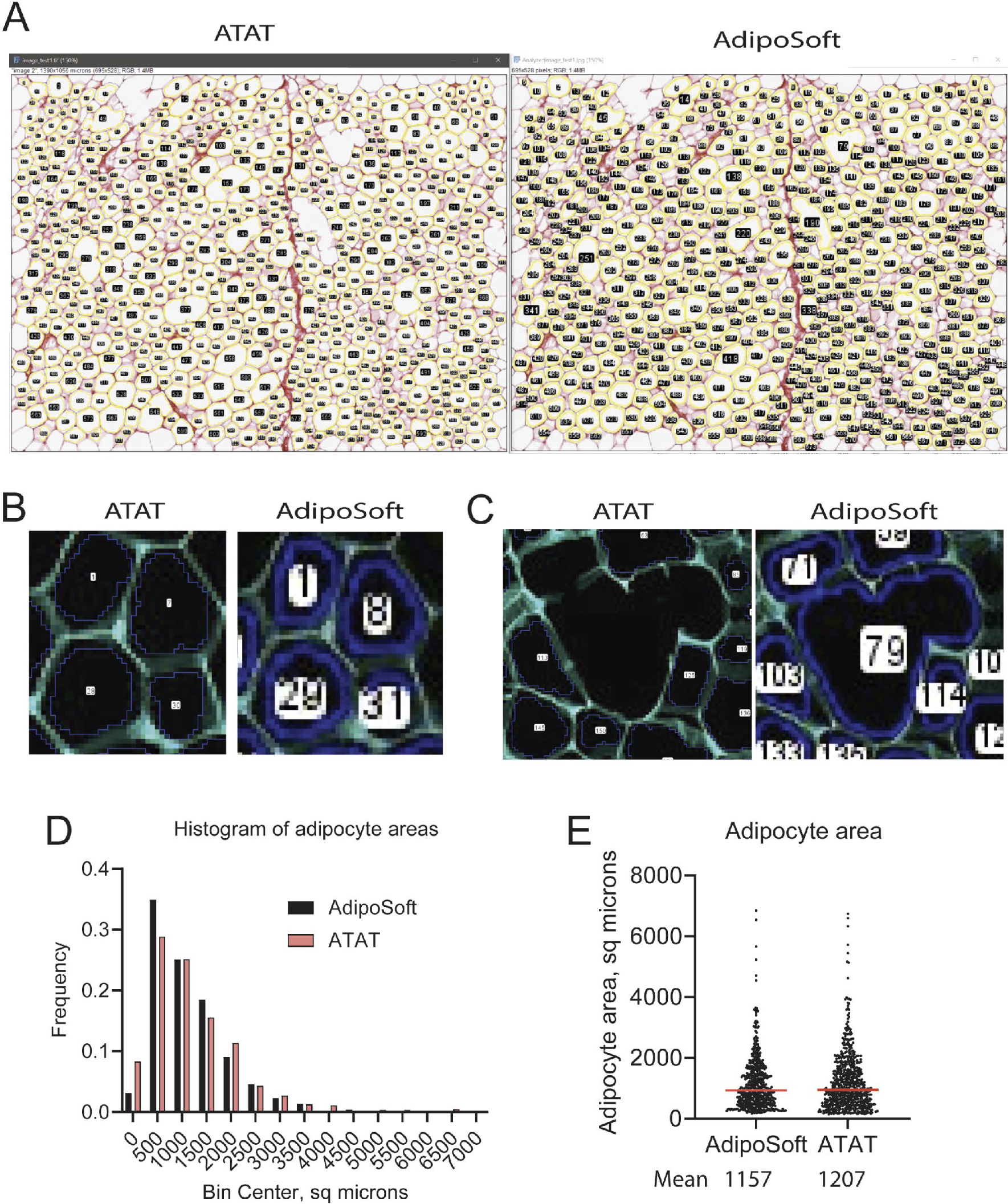
Comparative analysis of ATAT and Adiposoft. (A) Adipocyte capture by ATAT (left) and Adiposoft (right) in a representative image of macaque OM-WAT stained with Picro-Sirius Red. (B) A magnified view displaying the differences in adipocyte capture between ATAT and Adiposoft. (Note: Image is color-inverted for visualization purposes). (C) An example of an artifact originating from several adipocytes with ruptured membranes. ATAT excludes these artifacts from the analysis (Note: Image is color-inverted for visualization purposes). (D) Histogram illustrating the frequency distribution of adipocyte areas recorded by ATAT and Adiposoft. (E) Scatter plot depicting the comparison of adipocyte areas measured by ATAT and Adiposoft.

**Figure S2.**
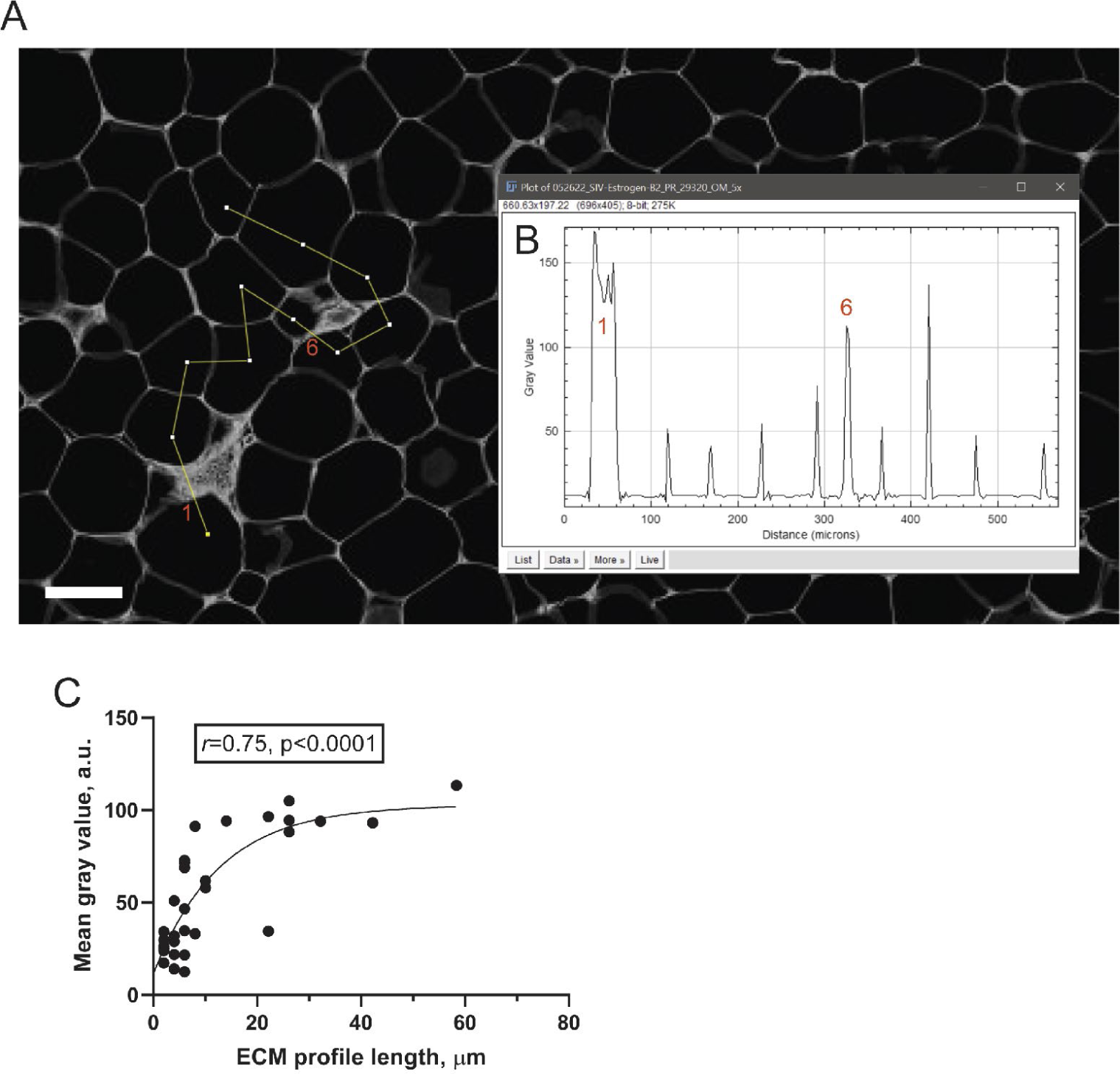
Correlation between ECM thickness and Picro-Sirius Red staining intensity. (A) Analysis of pericellular ECM profiles in a macaque OM-WAT section stained with Picro-Sirius Red. The yellow lines represent linear profiles drawn across pericellular ECM regions of multiple adipocytes; scale bar=100 microns. The analysis was conducted on a 8-bit inverted image, using a “plot profile” function in ImageJ. (B) Representative ECM profiles across 10 adipocytes, as shown in (A), illustrating the linear position in microns and pixel grey values. Thicker ECM regions are identified with numbers. (C) Correlation between the length of pericellular ECM and the mean grey pixel values of Picro-Sirius Red staining in pericellular ECM regions (n=31; 2 independent images). Pearson’s correlation coefficient (r) and p-value are indicated; the data is fitted with an exponential curve.

**Figure S3.**
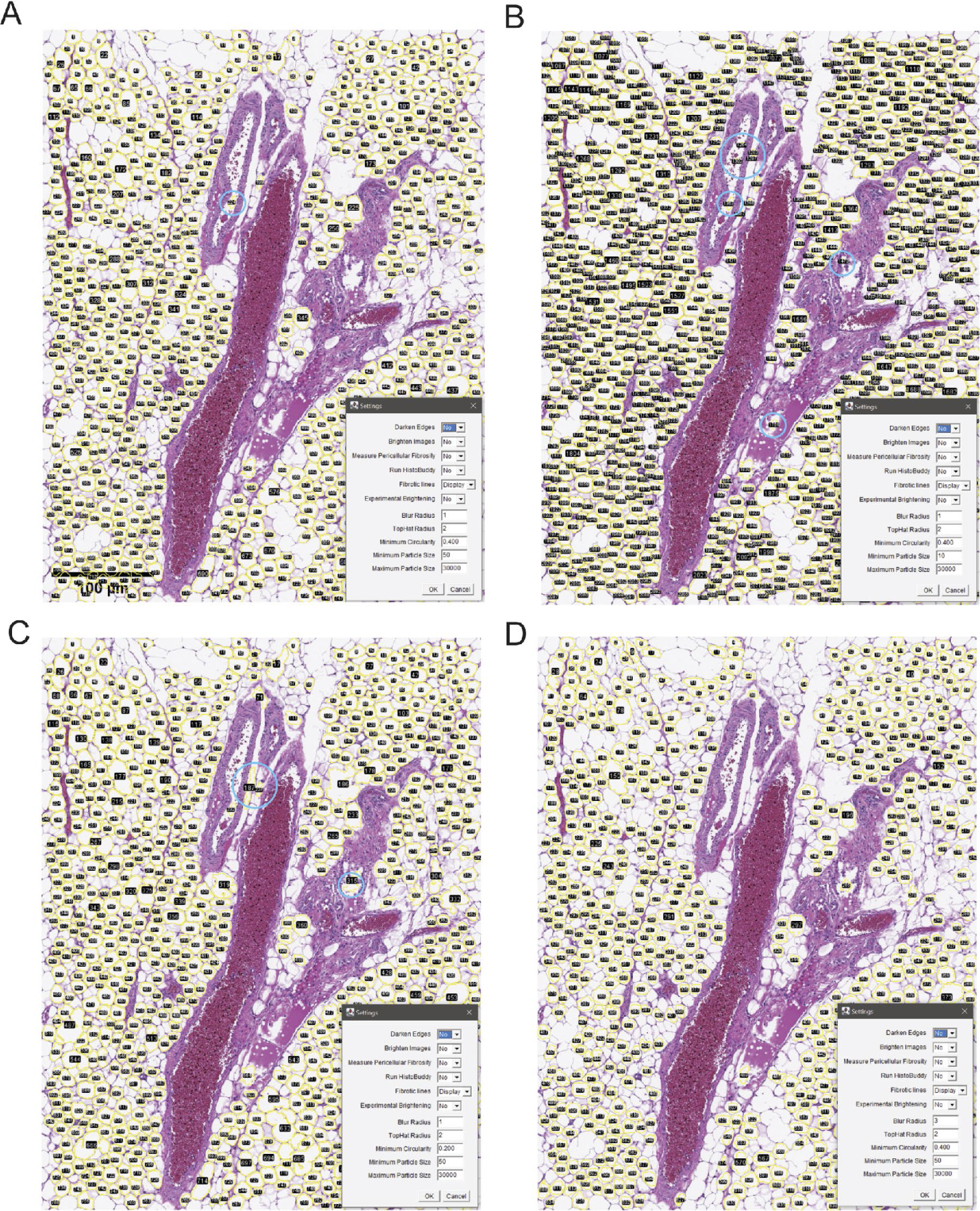
ATAT analysis of vascularized WAT areas. (A-D) Whole-image capture of adipocytes in H&E stained macaque OM-WAT, representing different capture parameters defined in “ATAT settings.” Decreasing particle size (A and B) and minimal circularity (A and C) increases adipocyte capture efficiency but may also increase the number of intravascular artifacts captured by ATAT. Increasing the blur radius (A and D) can decrease the number of some intravascular artifacts. User-defined parameters should be determined experimentally before analyzing large batches of images.

